# Retrotransposons hijack alt-EJ DNA repair process for mobilization and eccDNA biogenesis

**DOI:** 10.1101/2022.09.25.509424

**Authors:** Fu Yang, Weijia Su, Oliver W. Chung, Lauren Tracy, Lu Wang, Dale A. Ramsden, ZZ Zhao Zhang

## Abstract

Parasitic retrotransposons populate in the genome to rewrite the hosts’ genetic code. How retrotransposons use the host factors to achieve their propagation remains largely unclear. Here we report retrotransposons hijack the alternative end-joining (alt-EJ) DNA repair process to accomplish mobilization cycle. We applied Nanopore sequencing to examine the fates of replicated retrotransposon DNA, and found that only 10% of them achieve integration, while 90% exist as extrachromosomal circular DNA (eccDNA). Using eccDNA production as a readout, further genetic screens identified factors from alt-EJ as essential for retrotransposon replication. alt-EJ drives the 2^nd^-strand synthesis of the retrotransposon DNA via a circularization process, thus is necessitated for eccDNA production and integrations. Our study reveals a conserved function of alt-EJ in promoting selfish element propagation, which potentially causes disease or drives evolution.

---

Transposons abundantly occupy the genomes of nearly all animals, comprising almost 50% of human DNA (1-3). Given their ability to mobilize and (re)wire gene regulation, transposons bring one layer of genome dynamics to (re)writing genetic codes. Our previous efforts established *Drosophila* oogenesis as a platform to precisely characterize transposon activity at the mobilization level within an animal (4). We found that retrotransposons rarely mobilize in germline stem cells (4), which upon differentiation produce developing oocytes and supporting nurse cells (5). Instead, retrotransposons use nurse cells as factories to massively manufacture themselves like viruses (4). Then they transport the virus-like particles into the oocyte and mobilize into the genome that will be transmitted to the next generation (4). Leveraging this unique biological system that allows us to spatiotemporally follow their activation process, we sought to characterize how retrotransposons generate new copies of integrated DNA and (re)write the host germline genome.

For hundreds transposon families from the *Drosophila* genome, very few can achieve mobilization into oocytes (4). Among them, the LTR-retrotransposon *HMS-Beagle* displays the highest mobilization rate in oocytes (4). To thoroughly investigate its mobilization, we generated a fly strain carrying one copy of eGFP-tagged *HMS-Beagle* (Figure S1A). Landing it into a specific site of the fly genome, this eGFP-tagged *HMS-Beagle* serves as the sole precursor for any newly integrated copies that harbor eGFP sequences (Figure 1A and 1B). To potentially capture the *bona fide* mobilization events from this tagged *HMS-Beagle* within oocytes, we sequenced their genome using Nanopore technology, which can directly read DNA up to mega bases without PCR amplification.

**Figure 1.**
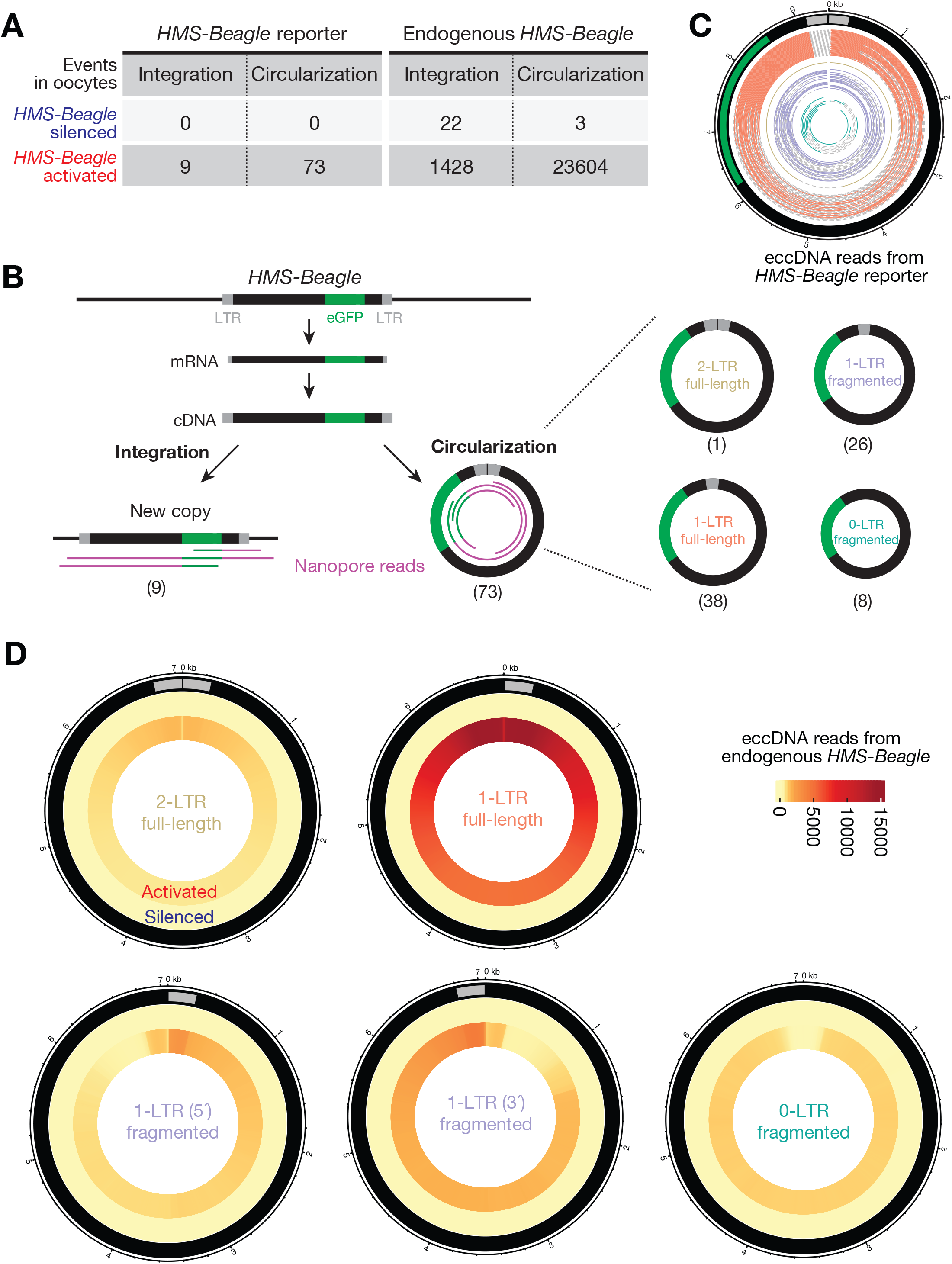
*HMS-Beagle* predominantly produces eccDNA upon activation. (A) Table to summarize the outcomes of replicated *HMS-Beagle* DNA detected in *Drosophila* oocytes. *HMS-Beagle* activation is achieved by suppressing Aub and Ago3 during oogenesis. (B) Workflow to characterize the integration and circularization (eccDNA) events from an engineered *HMS-Beagle* reporter. The Nanopore sequencing reads were classified as integration by having flanking sequences mapped to the genome or as eccDNA by containing end-to-end junction sites. The eccDNA was further classified into four categories based on their structures. The numbers within parentheses denote the number of reads identified for each type of event. (C) eccDNA reads from engineered *HMS-Beagle* reporter. Each circle represents a read: the solid part represents the sequenced region, and the dashed line represents the gap filled computationally. Salmon: reads supporting 1-LTR full-length circles; gold: one read supporting 2-LTR full-length circle; purple: reads supporting 1-LTR fragmented circles, likely resulting from autointegration; dark green: eccDNA reads do not contain intact LTR (D) Circos plot showing eccDNA reads from endogenous *HMS-Beagle*. Color scores indicate the mapping coverage throughout the full-length *HMS-Beagle* consensus. Outer layer: eccDNA reads from *HMS-Beagle* silenced oocytes. Inner layer: eccDNA reads from *HMS-Beagle* activated oocytes.

As expected, in the oocytes with transposons silenced (Figure S1B), the eGFP-tagged *HMS-Beagle* was only detected at its original landing site (Figure S1C and S1D).

We next triggered transposon activation by depleting Aub and Ago3 (Figure S1B), which are two key factors from a small RNA (piRNA) based silencing system that suppresses transposons during *Drosophila* oogenesis (6-8). Under this condition, we detected 9 new insertions from the tagged *HMS-Beagle* with 28X genome coverage (Figures 1A and S1D), consistent with our previous finding that *HMS-Beagle* preferentially targets the oocyte genome for integration (4).

Remarkably, manually analyzing the eGFP-derived reads that did not support integration indicated the formation of circular DNA. Since we prepared genomic libraries by the Tn5 tagmentation method, this would linearize any circular DNA molecules. Sequencing such molecules would allow us to quantify these DNA circles by searching for reads that cover the end-to-end junctions. By examining these events, we found that no circles formed when *HMS-Beagle* is silenced (Figure 1A). However, upon triggering its activation (Figure S1B), we observed 73 reads that support the production of circular DNA from eGFP-tagged *HMS-Beagle* with 28-fold genome coverage (Figure 1A and 1B), which is 8.1-fold more abundant than observed integration events. Given that *HMS-Beagle* has one LTR at each end, a head-to-tail circle would generate a junction read possessing two LTRs. However, among these 73 circle-supporting reads, only one has a junction with two LTRs (Figures 1B and 1C), suggesting that the formation of 2-LTR full length circles is a rare event. In contrast, 38 reads have junctions covering the start and end positions of the engineered *HMS-Beagle* reporter by containing one LTR (Figures 1B and 1C), indicating that 1-LTR full length circles are the dominant circular form. Among the remaining 34 reads that encompass fragmented *HMS-Beagle* sequences (termed as fragmented circles), 26 still have one intact LTR (termed as “1-LTR fragmented”). The remaining 8 reads do not contain intact LTRs (termed as “0-LTR fragmented”). Given their circular nature, we designated these *HMS-Beagle*-derived circles as extrachromosomal circular DNA (eccDNA). Collectively, our data suggest that upon activation, our engineered *HMS-Beagle* abundantly produced eccDNA as 1-LTR full length circles, but achieved far fewer integration events.

Our findings from the engineered *HMS-Beagle* prompted us to explore whether its endogenous copies also form circular DNA upon activation. We accordingly mined our Nanopore sequencing data to characterize the reads that support either integrated DNA or eccDNA events from endogenous *HMS-Beagle*. For control oocytes, in which transposons are silenced, we detected 0.8 potential integrations and 0.1 potential eccDNA from endogenous *HMS-Beagle* per genome coverage (Figures 1A and S2A).

These integration events most likely reflect polymorphisms between the genome of the fly strain used in this study and the *Drosophila* reference genome, hence defining the false-positive rate of our methodology on probing transposition events.

Upon transposon activation, we detected 1428 integrations and 23604 eccDNAs from endogenous *HMS-Beagle* loci with 28X genome coverage (Figures 1A, 1D, and S2A), highlighting that 94.3% of the replication products from *HMS-Beagle* form circles. Similar to the eGFP tagged reporter, endogenous *HMS-Beagle* appears to also primarily form 1-LTR full-length circles: 54.7% of eccDNA reads support 1-LTR full-length circles, 6.4% of eccDNA reads indicate 2-LTR full-length circles, and 38.9% of eccDNA reads are derived from fragmented *HMS-Beagle* circles (Figure 1D). We concluded that, consistent with the observations from our reporter, endogenous *HMS-Beagle* also dominantly forms 1-LTR eccDNA.

To validate the formation of eccDNA and the abundance of 1-LTR circles, we designed a set of divergent PCR primers, which would only give a PCR product upon the circularization of *HMS-Beagle* DNA (Figure 2A and S2C). Given that *HMS-Beagle* replication occurs within the ovary during oogenesis (4), we reasoned that their eccDNA is readily generated within ovaries before egg laying. By using ovary DNA as a template, performing PCR with divergent primers generated two products with distinct sizes (Figure S2C). Sanger sequencing revealed that the longer product with very low abundance contains two LTRs. In contrast, the shorter PCR product with high abundance contains only one LTR. Thus, our data consistently indicate that upon activation, endogenous *HMS-Beagle* preferentially generates eccDNA, especially in the form of 1-LTR circles.

**Figure 2.**
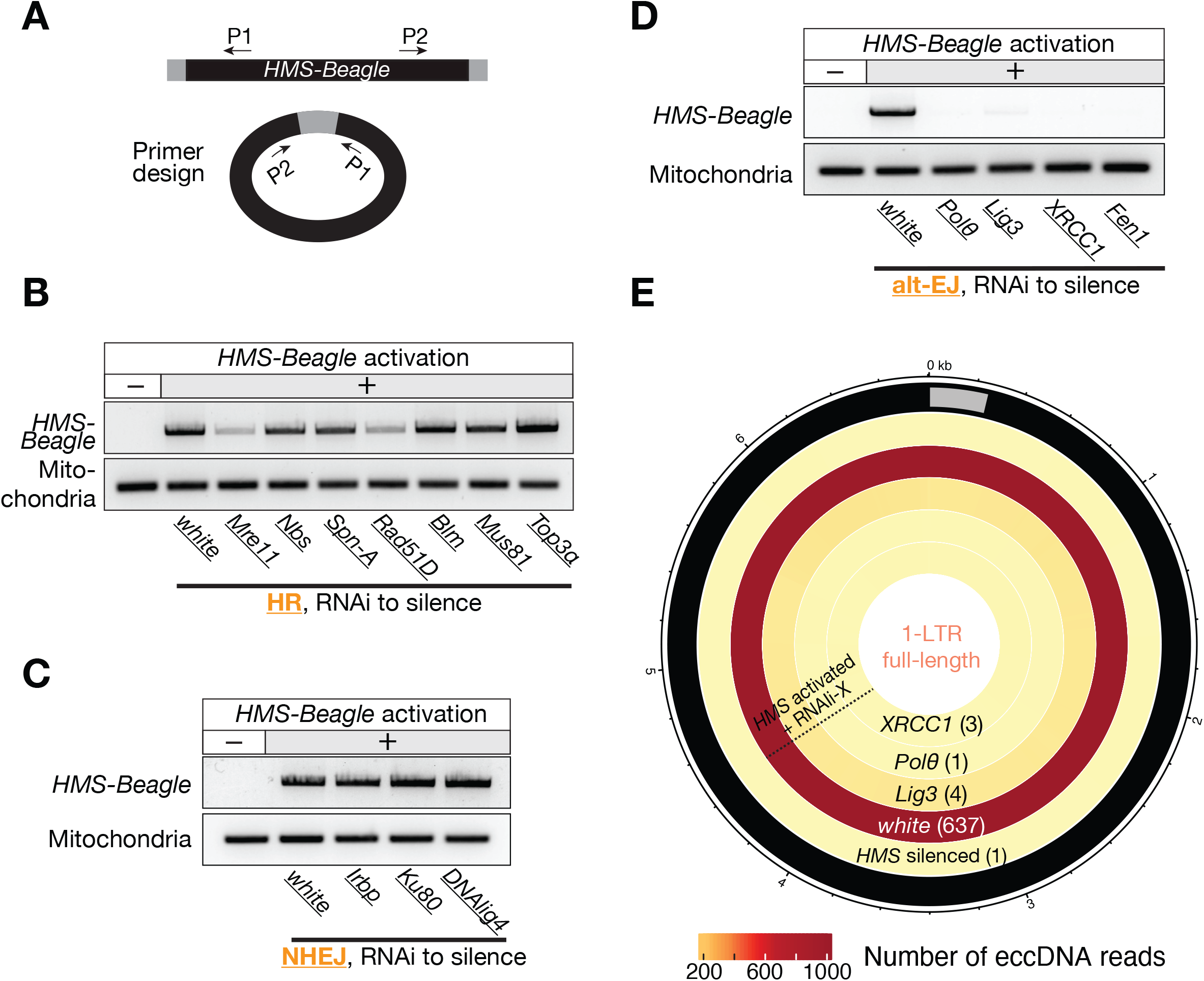
Factors from the Alt-EJ process drive 1-LTR full-length eccDNA formation. (A) Schematic of the design of divergent primers to identify *HMS-Beagle* eccDNA. (B-D) The representative gel image to show whether components from the HR (B), NHEJ (C), or Alt-EJ (D) process are required for the formation of 1-LTR full-length eccDNA. In this figure, transposon activation was achieved by silencing Aub in germline cells. (E) Circos plot showing 1-LTR full-length eccDNA reads from endogenous *HMS-Beagle*. Color scores indicate the mapping coverage throughout the full-length *HMS-Beagle* consensus. The numbers within parentheses denote the number of reads identified for each type of genotype.

Sequencing genomic DNA can indicate the formation of eccDNA by capturing the end-to-end junctions. However, this method lacks the power to validate their circularity or to reconstruct the full circular sequences of eccDNA. To obtain strong and direct evidence of circle formation, we established a Nanopore-based eccDNA sequencing method (eccDNA-Seq, Figure S3A). This method allows us to systematically characterize eccDNA production from *HMS-Beagle*. Applying eccDNA-Seq to normal fly ovaries generated reads mainly mapping to the mitochondrial genome (83.8% of total reads, Figure S3B), which is circular. This highlights the high efficiency of our method on enriching circular DNA for sequencing. In these control samples, *HMS-Beagle* generated very few, if any, eccDNA. By contrast, upon its activation, we detected 13362 *HMS-Beagle* eccDNA (Figure S3C). Consistent with our findings from genomic sequencing and PCR-based method, 62.47% of *HMS-Beagle* circles detected by eccDNA-Seq were 1-LTR full-length circles (Figure S3C). In summary, our eccDNA-Seq data provide strong and direct evidence of the formation of eccDNA from *HMS-Beagle in vivo*.

Retrotransposons use their RNA transcripts as a template for reverse transcription to generate DNA for subsequent integration (9). This replication intermediate could be the source for eccDNA biogenesis. This hypothesis predicts that depleting transposon RNA during oogenesis would abrogate eccDNA production. Indeed, once we depleted *HMS-Beagle* RNA by RNAi (Figure S4A), their eccDNA production was abolished (Figure S4B). Hence, our data indicate that retrotransposon-derived eccDNA is produced from their DNA replication intermediates, leaving their original genomic loci intact. This is different from the previously reported mechanisms on driving eccDNA formation, which involve either genomic DNA fragmentation or recombination within the genome (10-13).

For exogenous retroviruses, it has been proposed that the homologous recombination (HR) pathway can mediate the recombination of the two LTRs from the replicated linear DNA for the formation of 1-LTR circles (14, 15). To understand how *HMS-Beagle* retrotransposon forms 1-LTR eccDNA, we first tested the function of HR machinery proteins during this process. However, after individually depleting 7 key factors linked to this pathway during oogenesis––Nbs, Spn-A (*Drosophila* homolog of mammalian Rad51), Rad51D, Blm, Mre11, Mus81, and Top3a––*HMS-Beagle* still formed 1-LTR circles (Figure 2B). These data argue against a previously proposed function from the HR pathway in eccDNA biogenesis. Additionally, silencing of key factors from the non-homologous end joining (NHEJ) pathway (Irbp, Ku80, or DNA ligase 4) also had no impact on the formation of *HMS-Beagle*-derived eccDNA (Figure 2C).

To systematically characterize the factors that are essential for *HMS-Beagle* eccDNA formation, we performed a candidate-based RNAi screen to individually deplete 117 factors (from 129 alleles) that are known to function in DNA repair or DNA damage response (Table S2). After depleting each factor during oogenesis, we examined the production of *HMS-Beagle* 1-LTR circles by the PCR method we established (Figure S2C). Among these factors, 23 lead to lethality, impeding any further investigation (Table S2). From the rest of the candidates, four screened as essential factors for *HMS-Beagle* 1-LTR eccDNA production: DNA polymerase θ (Polθ, encoded by the *polQ* gene), XRCC1, DNA ligase 3 (Lig3), and Fen1 (Figures 2D, S5, and Table S2). Interestingly, all these four factors have been proposed to work coordinately for the alternative end-joining (alt-EJ) DNA repair process (also known as the microhomology-mediated end-joining, MMEJ) (16-18).

To further validate the function of alt-EJ factors on driving eccDNA production, we performed eccDNA-Seq upon individually depleting three of the identified factors: Polθ, XRCC1, and Lig3 (depleting Fen1 leads to semi-lethality, impeding obtaining enough DNA for sequencing). eccDNA-Seq generated consistent data with the PCR results: silencing each of these alt-EJ factors completely abolished the biogenesis of 1-LTR eccDNA from *HMS-Beagle* (Figure 2E).

How does alt-EJ process license eccDNA production from *HMS-Beagle*? It is possible that alt-EJ is required for transposon activation, and accordingly is necessitated for eccDNA production. To test this possibility, we performed Nanopore RNA-Seq and found the expression of *HMS-Beagle* transcripts is unaltered upon depletion of alt-EJ factors (Figure S6). This suggests that the alt-EJ factors are not required for retrotransposon transcription. Instead, here we provide evidence that alt-EJ is essential for the synthesis of *HMS-Beagle* 2^nd^-strand DNA via a circularization process, thus is necessitated for eccDNA biogenesis (Figure 3A).

**Figure 3.**
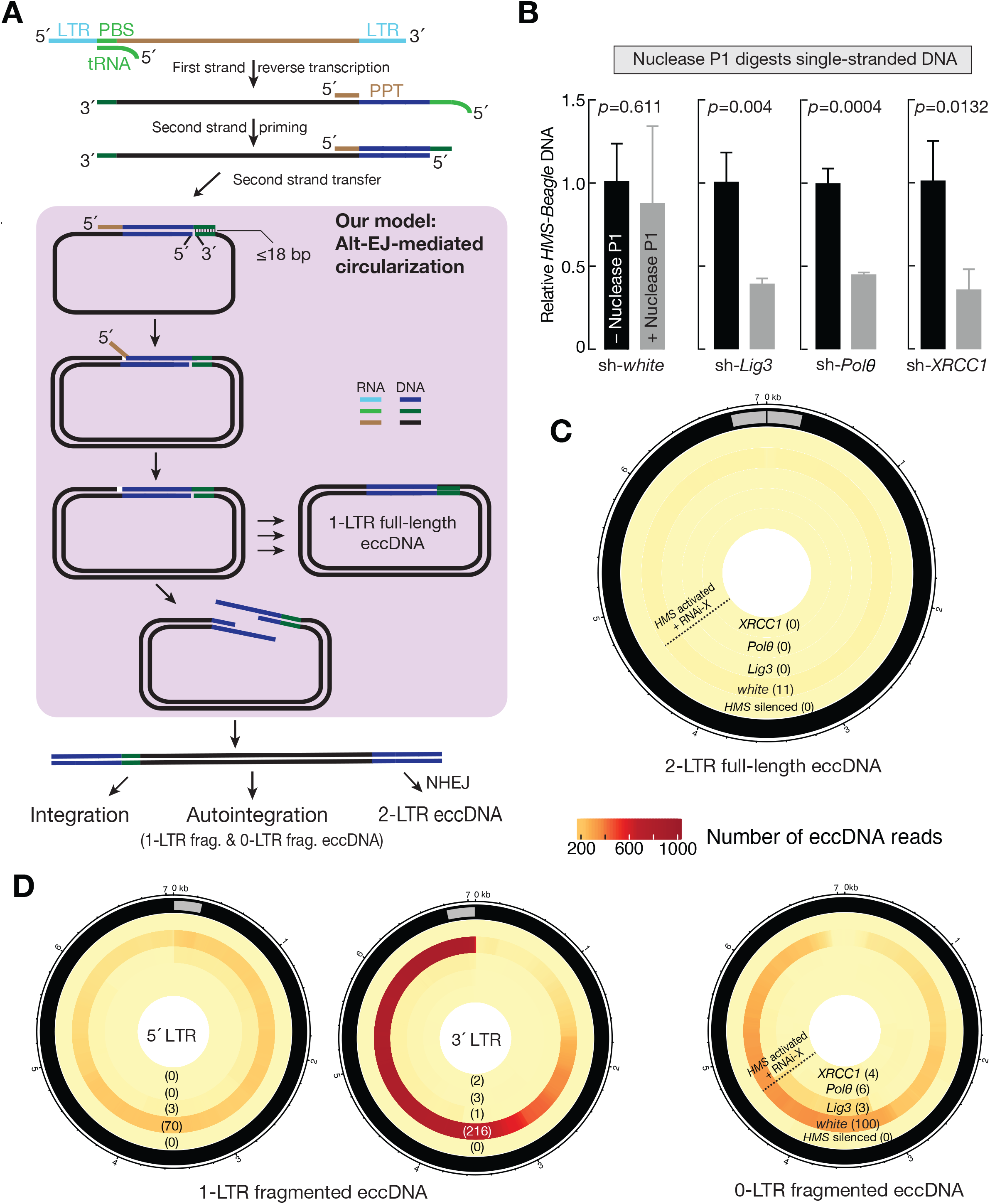
Blocking alt-EJ process abrogates DNA synthesis and all eccDNA production from *HMS-Beagle*. (A) A model to depict how alt-EJ-mediated circularization drives DNA synthesis, thus is essential for eccDNA production and mobilization. (B) qPCR to quantify the relative abundance of single-stranded *HMS-Beagle* DNA. All flies carry sh-*aub* to trigger transposon activation. The bars report mean ± standard deviation from three biological replicates. *p* values were calculated with a two-tailed, two-sample unequal variance *t* test. (C) Circos plot showing 2-LTR full-length eccDNA reads from endogenous *HMS-Beagle*. Color scores indicate the mapping coverage throughout the full-length *HMS-Beagle* consensus. The numbers within parentheses denote the number of reads identified for each genotype. These circles are likely generated by joining the 2 LTRs together via NHEJ. (D) Circos plot showing fragmented eccDNA reads from endogenous *HMS-Beagle*. Color scores indicate the mapping coverage throughout the full-length *HMS-Beagle* consensus. The numbers within parentheses denote the number of reads identified for each genotype. 1-LTR fragmented circles are likely generated by autointegration events: LTR to attack its own interstitial sequences *in cis*. 0-LTR fragmented circles are possibly the by-products of autointegration.

Like retroviruses, LTR-retrotransposons first transcribe genomic RNA (15). Using the 3’-end of a tRNA to pair with its primer binding site (PBS) sequence, the RNA transcripts can then convert to the 1^st^-strand DNA (15). Once finishing the 1^st^-strand DNA synthesis, the reverse transcriptase uses its RNase H activity to digest most of the retroviral RNA from the DNA-RNA hybrid except a short RNA sequence at the 3’ end– –the polypurine tract (PPT) (15, 19, 20). PPT serves as a primer to synthesize the 3’-LTR and the PBS sequence (by using the tRNA as template) for the 2^nd^-strand DNA (Figure 3A) (15, 20). To replicate the rest of the 2^nd^-strand sequence that is upstream of the PPT site, it has been proposed that the PBS sequences between the 1^st^ and 2^nd^-strand retroviral DNA can anneal with each other (15, 21). Known as the “2^nd^-strand transfer”, this step converts the 3’ end of the 2^nd^ strand DNA as the priming site for the synthesis of rest strand (15, 21). Despite the essentiality of this step during the life cycle of retrotransposons and retroviruses, what mediates this process remains mysterious.

Different from other DNA repair pathways, alt-EJ primes DNA synthesis by annealing a short homology (3-25 bp) (16, 18). Notably, the PBSs for retroviral elements are in general ≤ 18 nt (15, 22). Its apparent function in mediating microhomology formation led us hypothesize that alt-EJ circularizes the two DNA strands by annealing their PBS homology (Figure 3A). This circularization step initiates the subsequent 2^nd^-strand DNA synthesis. This would produce a non-covalent circle with two fates: either fill the nick to dominantly generate covalent 1-LTR eccDNA or convert into linear DNA with 2 LTRs (Figure 3A). The linear DNA can serve as precursors for three subsequent outcomes: forming 2-LTR circles, using its LTR to attack its own interstitial sequences *in cis* to generate fragmented circles (known as autointegration), inserting into host genomic DNA *in trans* for integration (Figure 3A). Our model predicts that impeding alt-EJ process would halt the 2^nd^-strand replication, thus leading to a higher single-stranded DNA ratio and the abolishment of all downstream outcomes. To rigorously test our model, we correspondingly quantified the production of single-stranded DNA (Figure 3B), 2-LTR eccDNA (Figure 3C), fragmented eccDNA (Figure 3D), and integration events upon the depletion of alt-EJ factors (Figure 4).

**Figure 4.**
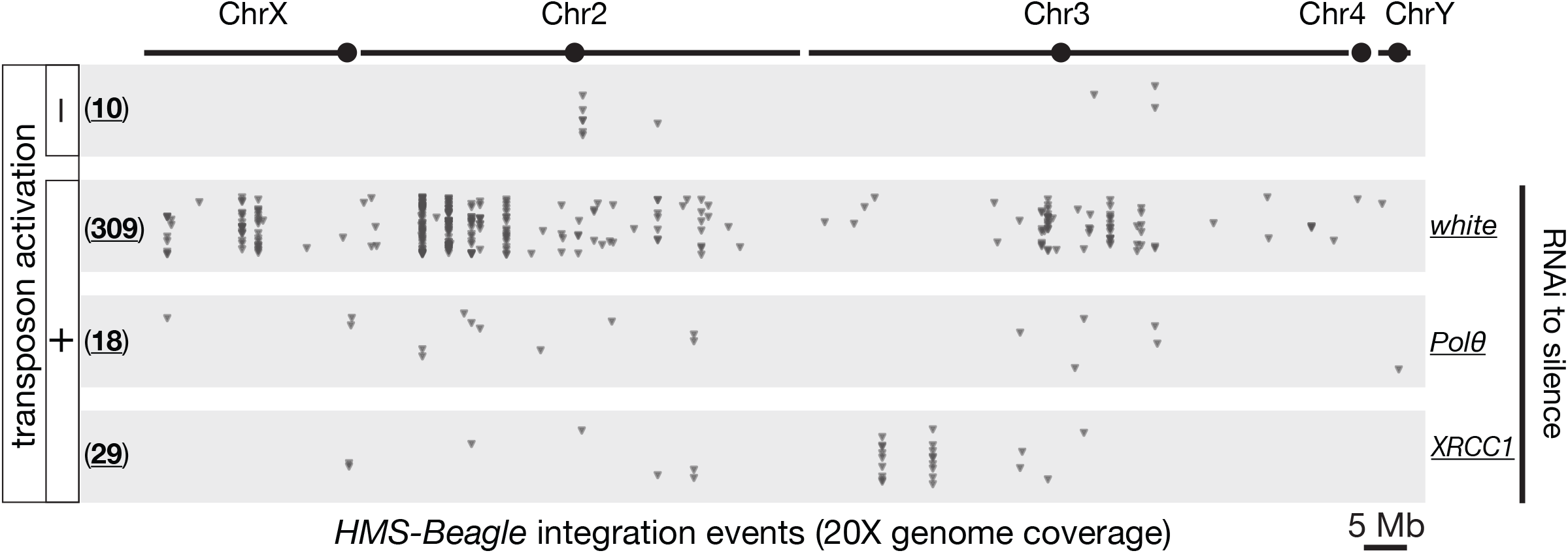
Blocking alt-EJ process abrogates *HMS-Beagle* mobilization. Dot plots to display the new integrations from *HMS-Beagle*. Transposon activation was achieved by silencing Aub in germline cells. Each triangle represents an integration event detected by Nanopore genome-seq. The numbers in parentheses present the total amounts of integration events detected. The numbers of integration detected under transposon-silenced condition likely represent the false-positive rates from our methodology.

First, we examined whether *HMS-Beagle* DNA extracted from the alt-EJ-perturbated ovaries is more sensitive to the treatment of Nuclease P1, an endonuclease that digests single-stranded DNA. Indeed, upon Nuclease P1 treatment, while *HMS-Beagle* DNA from control ovaries remained unchanged, silencing Polθ or Lig3 or XRCC1 led to > 2-fold reduction of *HMS-Beagle* DNA (Figure 3B). These data suggest that the alt-EJ process is essential for the completion of the 2^nd^-strand synthesis to produce double-stranded DNA.

Next, we detailedly examined the biogenesis of 2-LTR and fragmented eccDNA upon silencing of alt-EJ factors (Figure 3C and 3D). Here we report the eccDNA-Seq reads by normalizing to per mitochondrial genome coverage. For 2-LTR full-length circles, we detected 11 of them from *HMS-Beagle* after triggering its activation (Figure 3C). However, individually silencing Polθ or Lig3 or XRCC1 completely abolished their biogenesis (Figure 3C). Similarly, the number of fragmented circles also dropped to the background level upon suppressing the alt-EJ process (Figure 3D). Triggering *HMS-Beagle* activation generated 286 fragmented eccDNA that contain one intact LTR (1-LTR fragmented circles), which likely representing autointegration events. Meanwhile, there were 100 fragmented *HMS-Beagle* circles that do not contain intact LTRs (0-LTR fragmented circles, Figure 3D), which likely are generated as the by-products of autointegration events. However, for both categories of fragmented circles, we detected ≤ 6 circles with Polθ or Lig3 or XRCC1 silenced (Figure 3D). Our data thus further support a function of the alt-EJ pathway in the DNA replication process of LTR-retrotransposon.

Lastly, we tested whether impeding alt-EJ process would abrogate not only eccDNA production, but also *HMS-Beagle* mobilization. To test this, we individually depleted Polθ or XRCC1 (flies with Lig3 silenced lay very few mature oocytes), and then examined transposon mobilization rates in oocytes. For each genotype, DNA from somatic carcasses was sequenced to construct individual genomes, which serve as the reference to precisely define new transposon integrations in oocytes. With the same genome coverage (20X), we detected 309 new insertion events from *HMS-Beagle* when the alt-EJ process is undisturbed (Figure 4). However, once this process is blocked to abolish eccDNA biogenesis, the new insertion events also drastically decreased: to 18 insertions upon Polθ depletion, and 29 insertions upon XRCC1 depletion (Figure 4). Collectively, our data indicate that alt-EJ is essential for the generation of double-stranded *HMS-Beagle* DNA that serves as a precursor for both circularization and integration.

Besides using piRNA pathway perturbation to study *HMS-Beagle* eccDNA biogenesis and mobilization during oogenesis, we recently found that the *mdg4* retrotransposon, also known as *Gypsy*, naturally mobilizes in somatic tissues (*In press*). Particularly, *mdg4* appears to only mobilize at the pupal stage (*In press*), when flies are regenerating new somatic tissues for the adulthood. Accordingly, we monitored *mdg4* eccDNA production at different developmental stages. We found that mdg4 specifically generated eccDNA at the pupal stage, but not other developmental stages (Figure 5A and 5B), consistent with the time window when mobilization events are detected.

**Figure 5.**
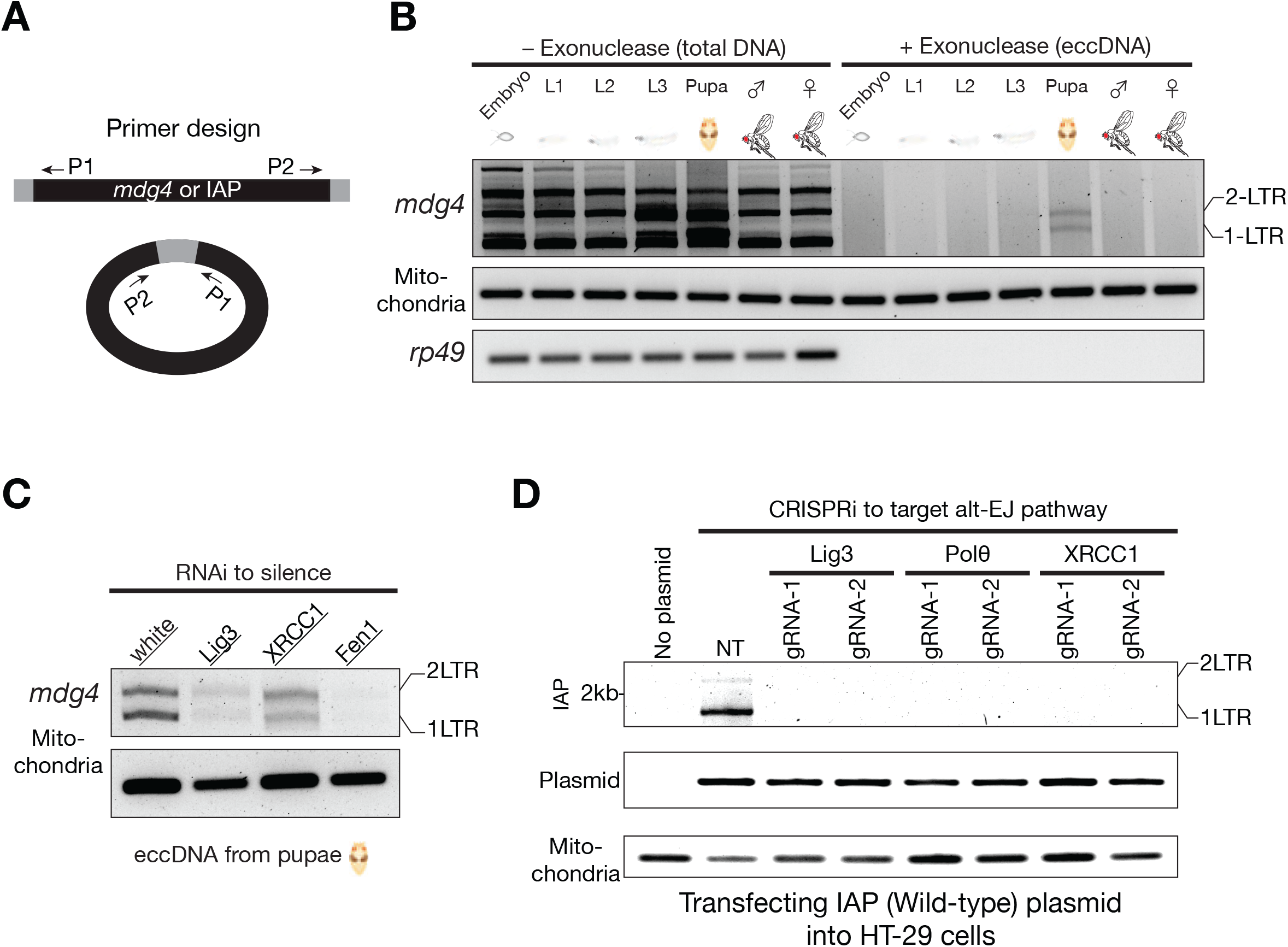
*mdg4* and mammalian IAP form eccDNA via the alt-EJ factors. (A) Schematic design of the divergent primers to detect *mdg4* or IAP eccDNA. (B) *mdg4* retrotransposon produces both 1-LTR and 2-LTR eccDNA at the pupal stage, the time window when mobilization occurs. Performing PCR using total DNA as template produced non-specific bands. Using exonuclease to enrich eccDNA generated two PCR products corresponding to 1-LTR and 2-LTR eccDNA respectively, as confirmed by Sanger sequencing. (C) eccDNA production from *mdg4* depends on alt-EJ factors. (D) Suppressing the factors from alt-EJ repair process blocks IAP eccDNA biogenesis. The PCR products corresponding to 1-LTR and 2-LTR eccDNA were confirmed by Sanger sequencing. NT (non-targeting) is a random gRNA without a targeting site.

Notably, silencing the alt-EJ factors suppressed *mdg4* eccDNA production (Figure 5C). These results suggest that the alt-EJ pathway is also essential for retrotransposon replication in somatic tissues.

Do mobile elements from different species also employ the alt-EJ process for their DNA replication? By using eccDNA production as a readout, we investigated the function of alt-EJ in the replication cycle of <u>I</u>ntracisternal <u>A</u>-<u>P</u>article (IAP), a mouse LTR-retrotransposon. IAP presents ∼2,800 full-length copies in the mouse genome and its activation contributes to ∼6% of all pathogenic mutations (23). To unambiguously examine IAP activity, previous work established a procedure to introduce IAP into cultured human cells (24, 25), which do not contain IAP in their own genome.

Following this procedure, we monitored IAP eccDNA formation in human cells and found that IAP indeed generated circular DNA, including both 1-LTR and 2-LTR circles (Figure 5D). Similar to *HMS-Beagle*, IAP dominantly generates 1-LTR circles (Figures 5D). Notably, upon individually depleting the human orthologs of the factors identified in *Drosophila* that function in the retrotransposon life cycle (Polθ, XRCC1, and Lig3; Figure S7), IAP failed to manufacture both 1-LTR and 2-LTR eccDNA (Figure 5D).

These findings indicate a conserved function of alt-EJ in driving the retrotransposon replication cycle in metazoan.

alt-EJ was initially viewed as a merely back-up pathway for canonical DNA repair (16, 18). By performing genetic screens *in vivo*, here we uncovered its conserved function in the replication cycle of parasitic genetic mobile elements. Expression of both alt-EJ factors and retrotransposons is tightly controlled. Notably, both of them appear to maintain a high activity during embryogenesis, or under aging or pathological progression, such as cancer (2, 16, 18, 26-30). Under these conditions, alt-EJ likely licenses eccDNA production and mobilization from retrotransposons, ultimately contributing to disease progression or driving evolution.

## AUTHOR CONTRIBUTIONS

Z.Z. and F.Y. conceived the project. All authors designed the experiments. For Figure 5D, L.T. designed primers and O.W.C. performed experiments. L.W. generated data for Figure 1. F.Y. performed all of the rest experiments. W.S. analyzed the sequencing data. Z.Z. wrote the manuscript. All authors read and approved the manuscript.

## ACKNOWLEDGEMENTS

We thank Marie Dewannieux and Kris Wood for providing plasmids, Julius Brennecke, Xin Chen, and BDSC for providing fly stocks. We thank members from ZZ lab for critical suggestions, and Don Fox, Xiao-Fan Wang, Bryan Cullen, Lin Lin, and David MacAlpine for reading the manuscript. This work was supported by the grants to Z.Z. from the Pew Biomedical Scholars Program and the National Institutes of Health (DP5 OD021355 and R01 GM141018), and to D.A.R from the National Cancer Institute (P01CA247773).

## DATA AVAILABILITY

The sequencing data were deposited to the National Center for Biotechnology Information (NCBI) with accession number PRJNA794176.

